# STING controls Herpes Simplex Virus *in vivo* independent of type I interferon induction

**DOI:** 10.1101/2019.12.12.874792

**Authors:** Lívia H. Yamashiro, Stephen C. Wilson, Huntly M. Morrison, Vasiliki Karalis, Jing-Yi J. Chung, Katherine J. Chen, Helen S. Bateup, Moriah L. Szpara, Angus Y. Lee, Jeffery S. Cox, Russell E. Vance

## Abstract

The Stimulator of Interferon Genes (STING) pathway initiates potent immune responses upon recognition of DNA derived from bacteria, viruses and tumors. To signal, the C-terminal tail (CTT) of STING recruits TBK1, a kinase that phosphorylates serine 365 (S365) in the CTT. Phospho-S365 acts as a docking site for IRF3, a transcription factor that is phosphorylated and activated by TBK1, leading to transcriptional induction of type I interferons (IFNs). IFNs are essential for antiviral immunity and are widely viewed as the primary output of STING signaling in mammals. However, other more evolutionarily ancestral responses, such as induction of NF-κB or autophagy, also occur downstream of STING. The relative importance of the various outputs of STING signaling during *in vivo* infections is unclear. Here we report that mice harboring a serine 365-to-alanine (S365A) point mutation in STING exhibit normal susceptibility to *Mycobacterium tuberculosis* infection but, unexpectedly, are resistant to Herpes Simplex Virus (HSV)-1, despite lacking STING-induced type I IFN responses. Likewise, we find *Irf3*^-/-^ mice exhibit resistance to HSV-1. By contrast, resistance to HSV-1 is abolished in mice lacking the STING CTT or TBK1, suggesting that STING protects against HSV-1 upon TBK1 recruitment by the STING CTT, independent of IRF3 or type I IFNs. Interestingly, we find that STING-induced autophagy is a TBK1-dependent IRF3-independent process that is conserved in the STING S365A mice, and autophagy has previously been shown to be required for resistance to HSV-1. We thus propose that autophagy and perhaps other ancestral interferon-independent functions of STING are required for STING-dependent antiviral responses *in vivo*.

## Introduction

The immune response to pathogens is initiated upon detection of pathogen-associated molecular patterns (PAMPs) such as lipopolysaccharide, flagellin and nucleic acids [1]. Double-stranded DNA (dsDNA) is an important PAMP for the detection of many pathogens, including *Mycobacterium tuberculosis* and Herpes Simplex Virus-1 (HSV-1) [2-4]. In vertebrates, the intracellular presence of dsDNA is detected by cyclic-GMP-AMP Synthase (cGAS), a dsDNA-activated enzyme that produces a cyclic dinucleotide (CDN) second messenger called 2′3′-cyclic-GMP-AMP (2′3′cGAMP) [5-10]. 2′3′cGAMP binds and activates the ER-resident transmembrane protein Stimulator of Interferon Genes (STING). Transcriptional induction of type I IFNs is widely presumed to be the primary output of STING signaling during antiviral defense. However, STING is evolutionarily ancient, present even in bacteria [11] and in animals such as the starlet sea anemone *Nematostella vectensis* and *Drosophila melanogaster* that do not appear to encode type I interferons [12]. By contrast, autophagy and NF-κB signaling are ancestral STING-induced signaling pathways, present in both *N. vectensis* and *D. melanogaster*, raising the possibility that these pathways are the primary or ancestral signaling outputs of STING [13-16].

The relative *in vivo* importance of the various signaling outputs of STING for anti-viral immunity in vertebrates is unknown. To address this issue, we used CRISPR/Cas9 to generate two distinct *Sting* mutant mouse lines: (1) STING S365A mice, which harbor a mutation in *Sting* that results in a serine to alanine substitution at amino acid 365; and (2) STING ΔCTT mice, in which valine 340 has been substituted by a STOP codon, resulting in a STING protein that lacks the entire CTT (Supp. Fig. S1a and S1b). We compared the S365A and ΔCTT mice to our previously generated STING-null *Goldenticket* (*Gt*) mice [17]. Since phosphorylation of S365 in the CTT of STING is required for the recruitment and activation of IRF3 [18-20], we predicted that S365A mice would be deficient in type I IFN responses downstream of STING, but would retain all other STING-dependent signaling events such as autophagy or NF-κB induction. The STING CTT contains S365 and is also essential for recruitment of TBK1 [21, 22]. Thus, we predicted that ΔCTT mice should also be deficient in all TBK1-dependent responses downstream of STING.

In the present study, we found that STING mutations do not affect susceptibility to *M.tuberculosis*, while control of HSV-1 infection requires the STING CTT but, unexpectedly, is largely independent of S365- or IRF3-induced type I IFNs. Control of HSV-1 also required TBK1, suggesting that STING protects against HSV-1 upon TBK1 recruitment by the STING CTT, independent of IRF3 or type I IFNs. We found that STING-induced autophagy is a TBK1-dependent IRF3-independent process that is conserved in the STING S365A mice. Thus, our data provide *in vivo* support for the idea that autophagy induction and perhaps other ancestral interferon-independent functions of STING may be preserved in vertebrates for host defense.

## Results

### Defective type I IFN induction in STING S365A and ΔCTT macrophages

Prior studies identified serine 365 of mouse STING (S366 in human STING) to be essential for STING-induced type I IFN expression in transfected or transduced cells *in vitro* [18-20]. To test whether endogenous STING requires the CTT and S365 for IFN induction in primary cells, bone marrow-derived macrophages from wild-type (WT) C57BL/6J, *Goldenticket* (*Gt*) STING null mice, and STING S365A and ΔCTT mice were stimulated with STING-specific agonists, including CDNs such as c-di-GMP and 2′3′cGAMP, as well as the cGAS agonist, dsDNA. As controls, cells were also stimulated with Sendai virus (SeV) and poly I:C, which induce type I IFNs via the RIG-I–MAVS pathway, independently of cGAS–STING. As expected, stimulation with STING-specific agonists resulted in increased *Ifnb* expression only in WT cells and not in any of the STING mutant cells. By contrast, the IFN response of all four genotypes was similar in response to SeV and poly I:C (Fig. 1a). STING activation can also lead to production of NF-κB-induced cytokines, such as TNF-α or IL-6 [23, 24]. Interestingly, primary *Gt*, S365A and ΔCTT macrophages stimulated *in vitro* with CDNs or dsDNA were defective for TNF-α induction as compared to WT cells (Supp. Fig. 1c). However, *in vivo* stimulation with 5,6-dimethylxanthenone-4-acetic acid (DMXAA), a potent STING agonist [25, 26], resulted in measurable TNF-α responses in the serum of WT and STING S365A mice, whereas *Gt* and ΔCTT mice were defective in TNF-α production as expected (Fig. 1b). As a control, the TNF-α response to STING-independent stimuli (e.g., LPS, which activates NF-κB via TLR4) was normal in all genotypes (Fig. 1b). We conclude that S365 may play a role in NF-κB activation, at least in macrophages, but is not required for NF-κB activation *in vivo* in response to strong STING agonists.

**Figure 1.**
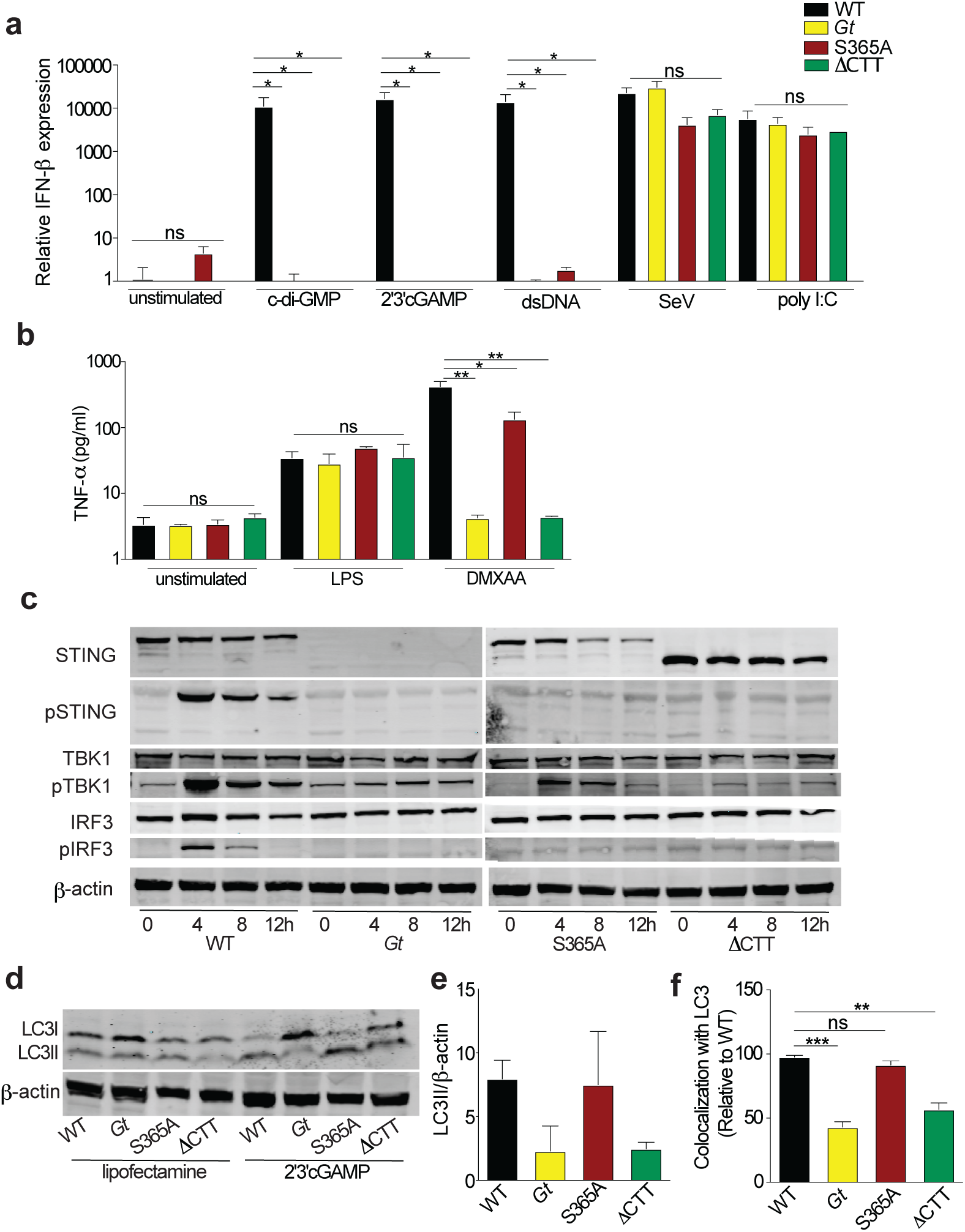
Defective type I IFN induction in STING S365A and ΔCTT macrophages. **a**, Bone marrow derived macrophages were stimulated for 6h and relative expression of IFN-β mRNA was measured. **b**, Mice were injected DMXAA (25 mg/kg, i.p.) or LPS (10 ng, i.v.) and TNF-α production was measured on the serum 2h later. **c**, Primary macrophages were transfected with dsDNA for 4, 8 or 12h or **d**, 2′3′cGAMP for 6h, and cell lysates were analyzed by immunoblotting for the indicated proteins. **e**, Quantification of (d). **f**, Quantification of LC3-DNA colocalization in primary macrophages transfected with Cy3-DNA for 6h and stained with LC3. Images were analyzed by an automated pipeline created on Perkin Elmer Harmony software for colocalization quantification (for more details refer to Methods). Representative results of three independent experiments. Error bars are SEM. Analyzed with one-way ANOVA and Tukey post-test. *, p ≤ 0.05; **, p ≤ 0.005; ***, p ≤ 0.0001. ns, not significant.

To further characterize our new STING mutant mice, the expression and/or activation of STING and downstream signaling components was assessed by immunoblotting (Fig. 1c). The STING S365A mutation did not affect expression of the STING protein itself or downstream components such as TBK1 and IRF3. STING ΔCTT mice harbor a STING protein of the expected (decreased) molecular weight. Phosphorylation of TBK1—but not of STING or IRF3—occurred in S365A cells in response to STING agonist, consistent with the generally accepted requirement for S365 phosphorylation for IRF3 binding and activation (Fig. 1c). By contrast, no phosphorylation of STING, TBK1 or IRF3 was seen in ΔCTT cells, as expected.

In addition to its role in IFN-induction, TBK1 has previously been shown to activate autophagy via the phosphorylation of autophagy adaptor proteins such as NDP52, p62 and optineurin [27]. Likewise, STING activation itself is associated with autophagy-like responses [16, 19, 28, 29]. Interestingly, a recent report claimed that STING-induced autophagy does not require the CTT or TBK1 [16]; however, these experiments utilized conditions that may not reflect the true *in vivo* requirements, such as overexpressed proteins, immortalized cell lines, and/or artificial *in vitro* stimulations. In order to investigate whether S365 or the CTT is required for endogenous STING to activate autophagy-like processes, primary macrophages were transfected with 2′3′cGAMP and conversion of LC3-B from form I to the lipidated form II was analyzed. Robust LC3-B conversion was observed in WT and S365A cells, while this response was reduced in *Gt* and ΔCTT cells (Fig. 1d and e, Supp. Fig. S1d), indicating that STING-dependent autophagy is independent of S365A-IRF3 activation and type I IFN responses but largely requires the CTT. To confirm this result, we quantified colocalization of LC3 puncta and cytosolic DNA. Primary macrophages were transfected for 6h with Cy3-labeled DNA and colocalization with LC3 puncta was quantified by immunofluorescence. STING-deficient *Gt* and ΔCTT cells exhibited poor colocalization of DNA and LC3, whereas WT and S365A cells exhibited robust and indistinguishable DNA–LC3 colocalization (Fig. 1f and Supp. Fig. S1e). Taken together these data indicate that endogenous STING requires S365 for IRF3 recruitment and induction of type I IFNs downstream of STING, whereas the CTT (but not S365) is required for TBK1 recruitment and robust autophagy induction. Our new mouse models therefore allow us to genetically separate the IFN- and autophagy-inducing functions of endogenous STING for the first time.

### STING mutant mice exhibit normal susceptibility to *M. tuberculosis* infection

To determine whether STING-induced interferons and autophagy have distinct functions during *in vivo* infection, we first examined infections with the bacterium *M. tuberculosis*. Previous reports have suggested that the cGAS–STING pathway detects *M. tuberculosis* in macrophages and initiates both a type I IFN response and autophagy-like colocalization of bacteria with LC3 [4, 30-32]. Type I IFNs exacerbate many bacterial infections, including *M. tuberculosis* infection [33-36] whereas autophagy is generally anti-bacterial [37]. Therefore, loss of STING may have counteracting effects that obscure its function; indeed, STING-null *Gt* mice do not exhibit dramatic alterations in susceptibility to *M. tuberculosis* infection [31, 38]. We hypothesized that perhaps STING S365A mice, which are defective for STING-induced type I IFN induction but not for autophagy, might exhibit enhanced resistance to *M. tuberculosis*. Consistent with this hypothesis, *Irf3*^-/-^ mice have previously been reported to be resistant to *M.tuberculosis* [30]. Therefore, we aerosol infected mice harboring WT, *Gt*, S365A, or ΔCTT STING alleles with virulent *M.tuberculosis*. We found that all STING genotypes were similarly susceptible to *M.tuberculosis* with similar survival rates, bacterial burdens in lungs and spleens, and cytokine production (Fig. 2a-l).

**Fig 2.**
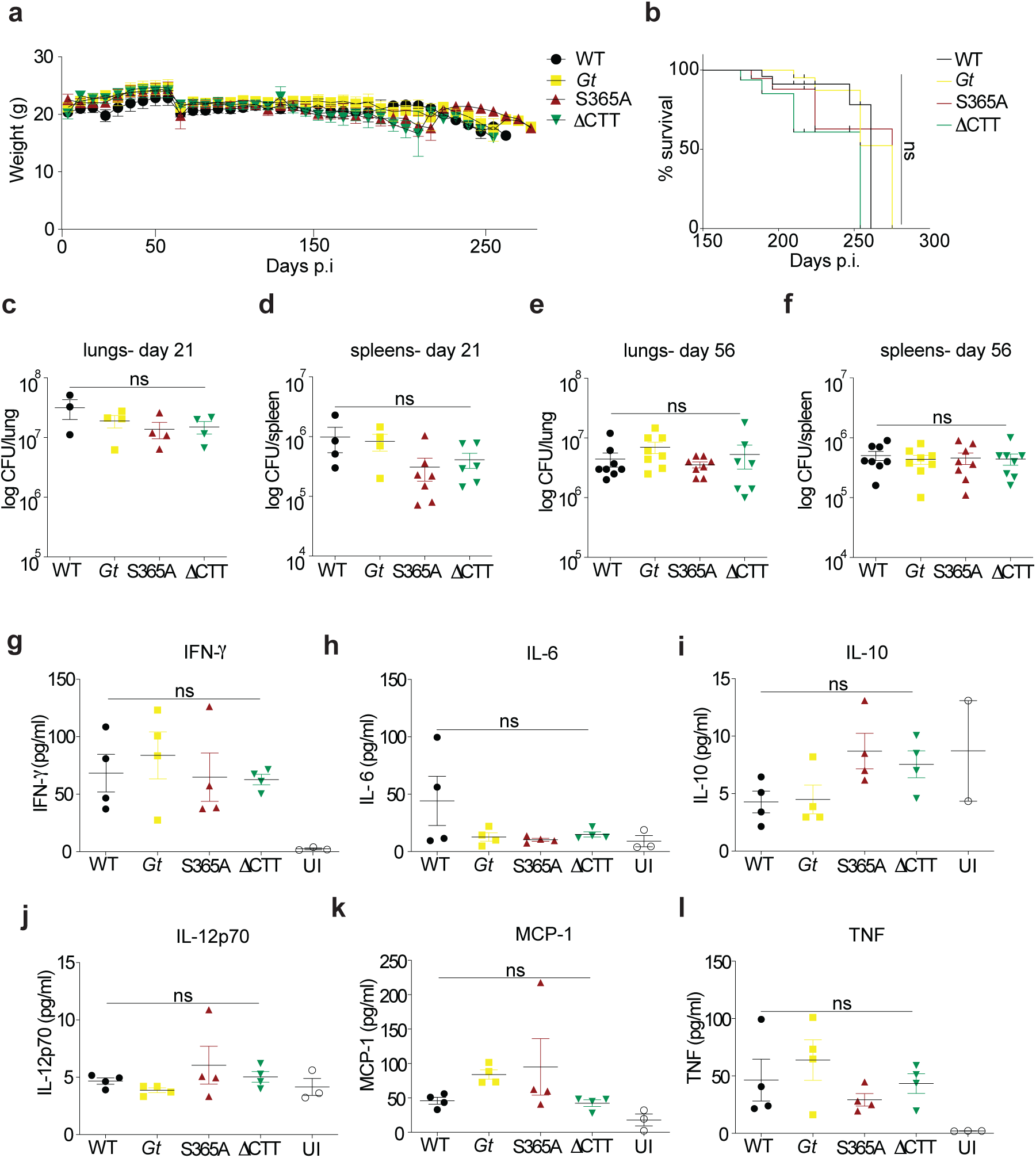
STING mutant mice exhibit normal susceptibility to *M.tuberculosis* infection. Mice were aerosol infected with 400 CFU dose of *M. tuberculosis* (Erdman strain) and **a**, weighed every week. **b**, Survival of infected mice. **c**, Bacterial burden from lungs and **d**, spleens at 21 and **e-f**, 56 days post infection. **g-l**, Cytokine levels in the lungs from infected mice at day 21, measured by CBA. Similar (not statistically different among genotypes) results were observed at day 56 (data not shown). All mice except C57BL/6J WT were bred in-house. Representative results of five independent experiments. Error bars are SEM. Analyzed with one-way ANOVA and Tukey post-test. ns, not significant.

To confirm that STING-induced type I IFN signaling does not affect *M. tuberculosis* susceptibility, we also sought to infect mice lacking the downstream transcription factor, IRF3. However, the published *Irf3*^-/-^ mice that were previously tested are also deficient in *Bcl2l12*, a gene that neighbors *Irf3* and that was inadvertently disrupted by the deletion targeting *Irf3* [39]. Therefore, we generated new *Irf3* deficient (but *Bcl2l12*^+/+^) mice, as well as *Bcl2l12*^-/-^ (but *Irf3*^+/+^) mice, using CRISPR–Cas9 (Supp. Fig. S2a-S2d). We found *Irf3*^-/-^ mice, *Bcl2l12*^-/-^ mice, and the previously tested doubly deficient mice, were all similarly susceptible to *M.tuberculosis* as WT mice (Supp. Fig. S2e-f). We cannot explain the previously reported resistance of *Irf3*^-/-^ mice but suspect this may be related to microbiota differences between *Irf3*^-/-^ lines. Nevertheless, we conclude that although *M. tuberculosis* can activate cGAS–STING– IRF3 in macrophages *in vitro*, STING does not appear to play significant beneficial or detrimental roles in *M.tuberculosis* pathogenesis *in vivo*.

### S365A mice are resistant to systemic HSV-1 infection

Given that STING is essential for resistance to HSV-1, we next decided to challenge our STING mutant mice with HSV-1. *Sting*-deficient mice are highly susceptible to HSV-1 infection [40-42]. Although induction of type I IFN is presumed to be a major mechanism of STING-mediated protection against HSV-1, the relative importance of type I IFNs and other STING-dependent responses in host defense against HSV-1 has not been resolved. Indeed, the immune response to HSV-1 is complex and multi-factorial. HSV-1 encodes factors to block the type I IFN response, perhaps limiting its effectiveness in control of the infection [41, 43, 44]. Moreover, it has been shown that neurons do not require type I IFNs—and can instead rely on autophagy—to limit HSV-1 replication in mice *in vivo* and *in vitro* [45]. These observations led us to hypothesize that interferon-independent signaling downstream of STING may contribute to control of HSV-1.

Initially, mice were intravenously infected with HSV-1 (KOS strain). As expected, WT mice were resistant to infection and remained healthy through 12 days post infection, whereas STING-deficient *Gt* mice were very susceptible to infection and exhibited rapid weight loss and complete paralysis, succumbing 6 days post infection (Fig. 3a–c) [41]. The ΔCTT mice phenocopied the susceptibility of *Gt* mice, demonstrating that the STING CTT is critical for defense against HSV-1. However, in contrast to ΔCTT mice, the S365A mice unexpectedly showed marked resistance to infection, exhibiting only limited weight loss and paralysis, and recovering fully after 6 days of infection (Fig. 3a–c). Susceptibility of *Gt* and ΔCTT mice correlated with elevated viral titers in the brains and spinal cords compared to reduced titers in resistant WT and S365A tissues (Fig. 3d and e). Viral titers among all four genotypes were similarly low in the liver, confirming the neurotropism of HSV-1 (Supp. Fig. S3a). Given that type I IFNs are essential for resistance to HSV-1 [46, 47], and that STING is required for type I IFN induction to HSV-1 [40-42, 48], we were surprised that S365A mice were not as susceptible to infection as *Gt* and ΔCTT mice. One possibility to explain this result is that S365A is not required for STING-dependent type I IFN induction *in vivo*. To test this possibility, we measured expression of *Ifnb* and the interferon stimulated genes (ISGs) *viperin* and *Ifit1* in mice brains following intravenous infection. Only WT brains exhibited a detectable STING-induced IFN response (Fig. 3f and g and Supp. Fig. S3b). In addition, *Tnf* and *Il6* expression was also elevated only in the brains of WT mice (Supp. Fig. S3c and S3d). These data indicate that S365 is critical for STING-induced type I IFN and other cytokines, but surprisingly, this S365-induced response is not critical for STING-dependent immunity to HSV-1.

**Figure 3.**
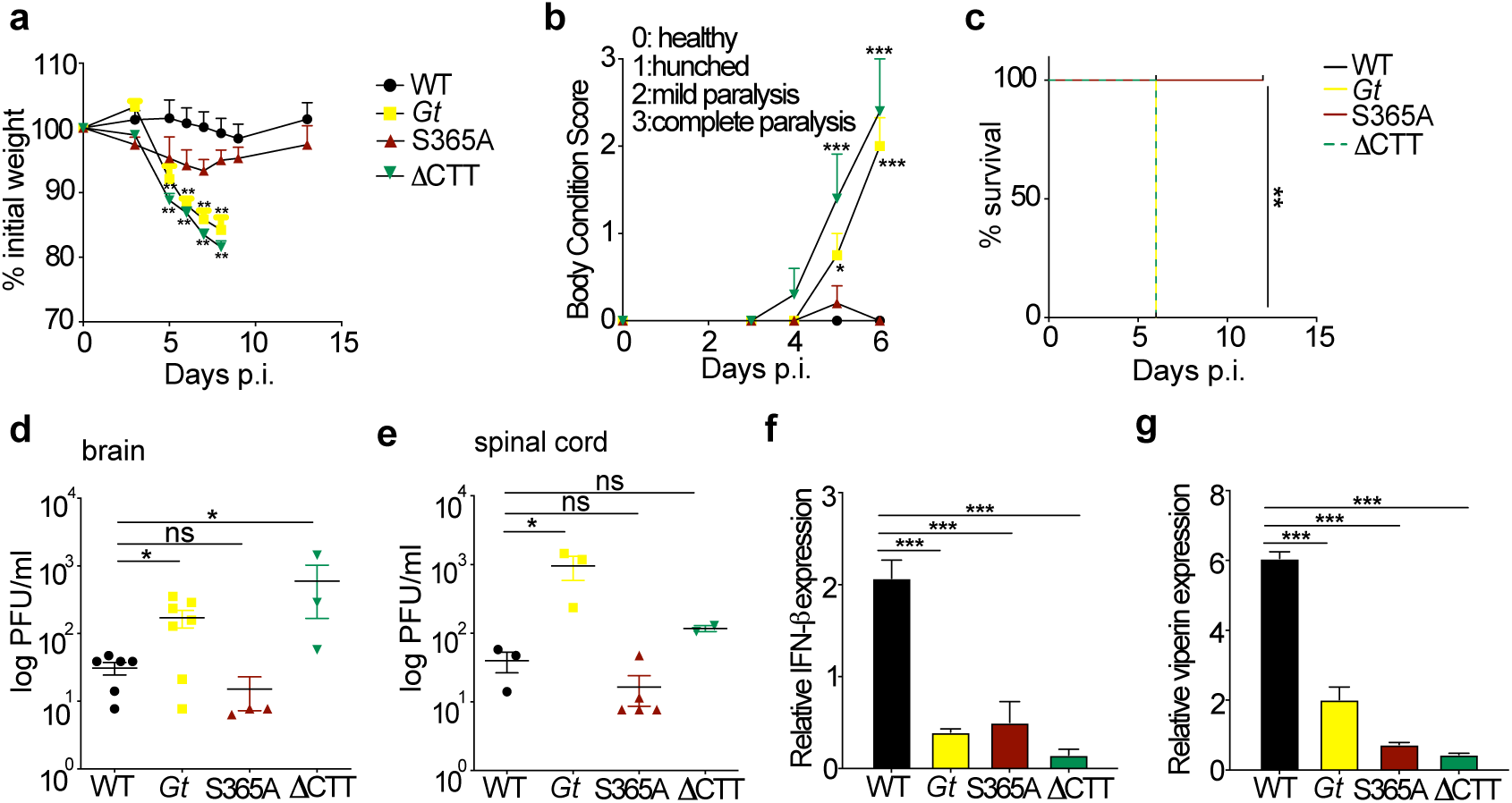
S365A mice are resistant to systemic HSV-1 infection. Mice were intravenously infected with 1×10^6^ PFU of HSV-1 (KOS strain). **a**, Percentage of initial weight following infection. **b**, Body condition score (BCS) of infected mice. **c**, Survival of mice following infection. **d**, Viral titers in the brain and **e**, spinal cord at 6 days p.i. **f**, Relative expression of *Ifnb* and **g**, *Viperin* in the brain at 3 days p.i. All mice except C57BL/6J WT were bred in-house. Representative of at least three independent experiments. Error bars are SEM. Analyzed with one-way ANOVA and Tukey post-test. *, p ≤ 0.05; **, p ≤ 0.005; ***, p ≤ 0.0001. ns, not significant.

### S365A mice are resistant to ocular HSV-1 infection

HSV-1 is a neurotropic virus that is transmitted via mucosal routes (typically oral, ocular, or genital) and infects epithelial cells before reaching the central nervous system where it establishes latency in neurons [49, 50]. Therefore, in order to mimic a more natural route of infection, we challenged mice with HSV-1 using an eye infection model [41, 51]. In these experiments, we used strain 17, a more virulent HSV-1 isolate, because the KOS strain used for intravenous infections fails to cause pathology in the eye infection model [51]. As with systemic infection, *Gt* and ΔCTT mice rapidly lost weight and all mice succumbed to infection by 6-7 days post infection (Fig. 4a, b). In contrast, WT mice remained fully resistant and S365A mice presented an intermediate phenotype, with initial weight loss but later recovery and ∼50% survival (Fig. 4a, b). Similar to systemic infection, the susceptibility of the mice correlated with viral burdens: WT and S365A exhibited lower viral titers in the eye wash (Supp. Fig. S4a), whole brain, brain stem and spinal cord as compared to *Gt* and ΔCTT mice (Fig. 4c-e). Once again, we found that *Ifnb* and *viperin* expression was elevated in WT but not in *Gt*, S365A or ΔCTT brain stems (Fig. 4f and Supp. Fig. S4b). Previous studies have shown that STING-dependent control of HSV-1 is cell-type specific [41]. To investigate an S365-dependent viral control in brain cells, we infected primary neurons and astrocytes *in vitro* with HSV-1. However, we observed similar viral yields and autophagy-related processes (colocalization of virus-LC3 and LC3 conversion) (Supp. Fig.S5a-e) in both cell populations among all genotypes, confirming prior reports that STING does not function cell autonomously in these cell types [42]. To address which cells require S365 for type I IFN induction *in vivo*, we sorted brain cells (neurons, astrocytes and microglia) 3 days post infection from brains of HSV-1-infected mice (ocular route). We found elevated *Ifnb* expression in all cell populations only in WT mice (Fig. 4g-i), confirming that IFN-β induction *in vivo* requires STING S365. Together, our data suggest that STING-mediated control of HSV-1 infection *in vivo* does not require STING S365-induced type I IFN production.

**Figure 4.**
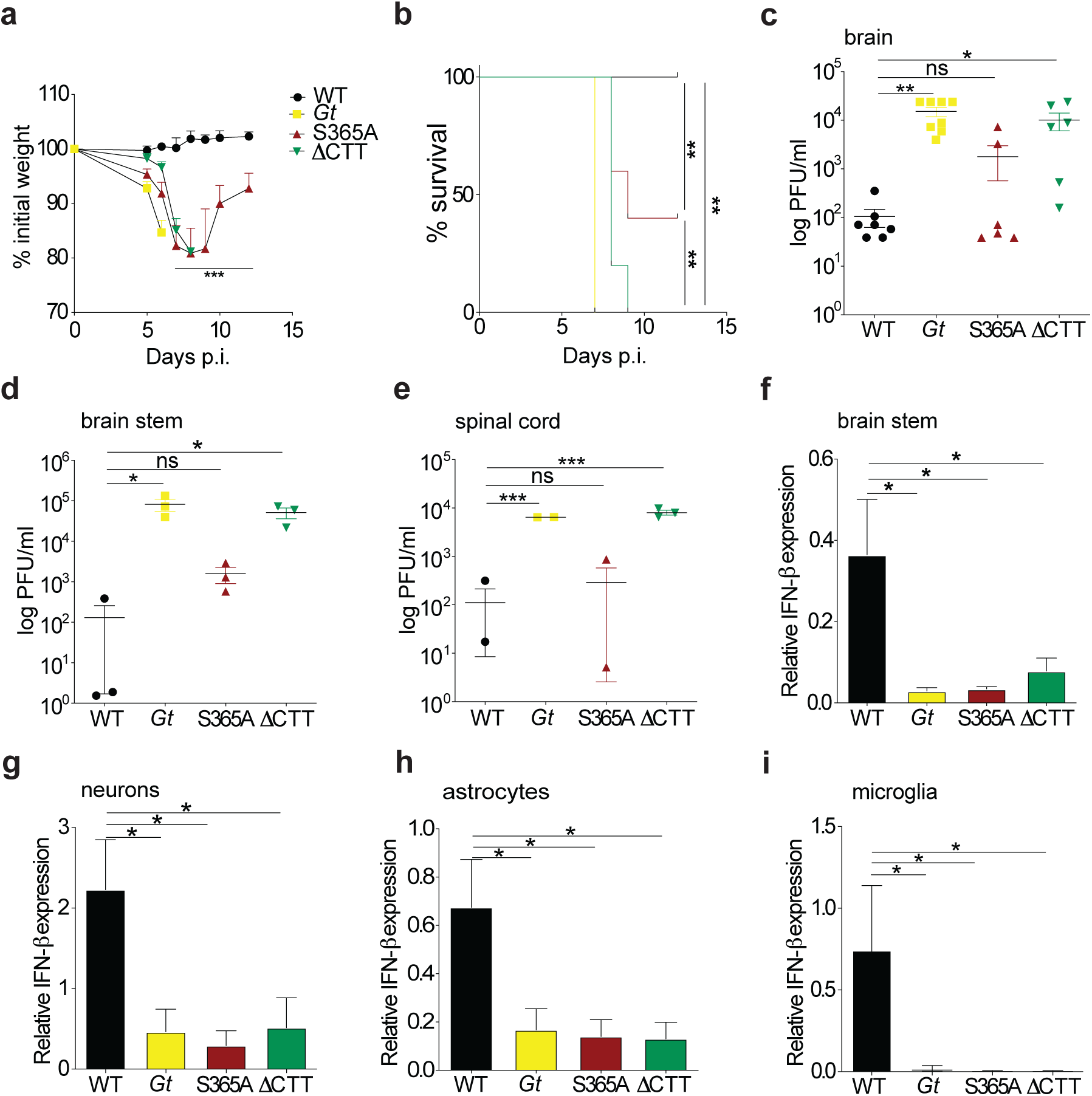
S365A mice are resistant to ocular HSV-1 infection. Mice were infected via the ocular route with 1×10^5^ PFU of HSV-1 (strain 17). **a**, Percentage of initial weight following infection. **b**, Survival of infected mice. **c**, Viral titers in the brain, **d**, brain stem and **e**, spinal cord from 6 days p.i. **f**, Relative expression of *Ifnb* in the brain stem at 3 days p.i. **g**, Brains from infected mice were collected 3 days p.i. and neurons, **h**, astrocytes and **i**, microglia cells were sorted, and *Ifnb* expression was analyzed. Representative of more than five independent experiments. Error bars are SEM. Analyzed with one-way ANOVA and Tukey post-test. *, p ≤ 0.05; **, p ≤ 0.005; ***, p ≤ 0.0001. ns, not significant.

### STING S365A and *Irf3*^*-/-*^ mice phenocopy resistance to HSV-1

Because TBK1 has been implicated in autophagy induction [4, 52, 53] whereas IRF3 acts as a transcriptional factor to induce type I IFNs downstream of STING, we investigated the role of these proteins in the context of an *in vivo* infection with HSV-1. *Tbk1*^-/-^ mice die as embryos, but this lethality is reversed on a *Tnfr1*^-/-^ background. We therefore analyzed *Tnfr1*^-/-^ mice compared to *Tbk1*^-/-^*Tnfr1*^-/-^ double deficient mice. *Tbk1*^-/-^*Tnfr1*^-/-^ mice lost weight and succumbed to HSV-1 infection at the same rate as *Gt* and ΔCTT mice, whereas *Tnfr1*^-/-^ mice were as resistant to HSV-1 as WT mice (Fig. 5a and b). By contrast, *Irf3*^-/-^ mice presented an intermediate phenotype similar to that of S365A mice. Viral loads in brain stems and total brain correlated with the disease severity (Fig. 5c and Supp. Fig. S6a) and *Ifnb* expression in the brain stems was increased only in WT mice (Fig. 5d). *Ifnb* expression was also reproducibly decreased in *Tnfr1*^-/-^ mice *in vivo* (but not *in vitro* Supp. Fig. S6b) for reasons that are currently unclear. Nevertheless, these results suggest that S365A is critical for STING-induced IRF3 activation and *Ifnb* expression, but neither S365 nor IRF3 are essential for restriction of HSV-1 replication *in vivo*, whereas the STING CTT and TBK1 are essential.

**Figure 5.**
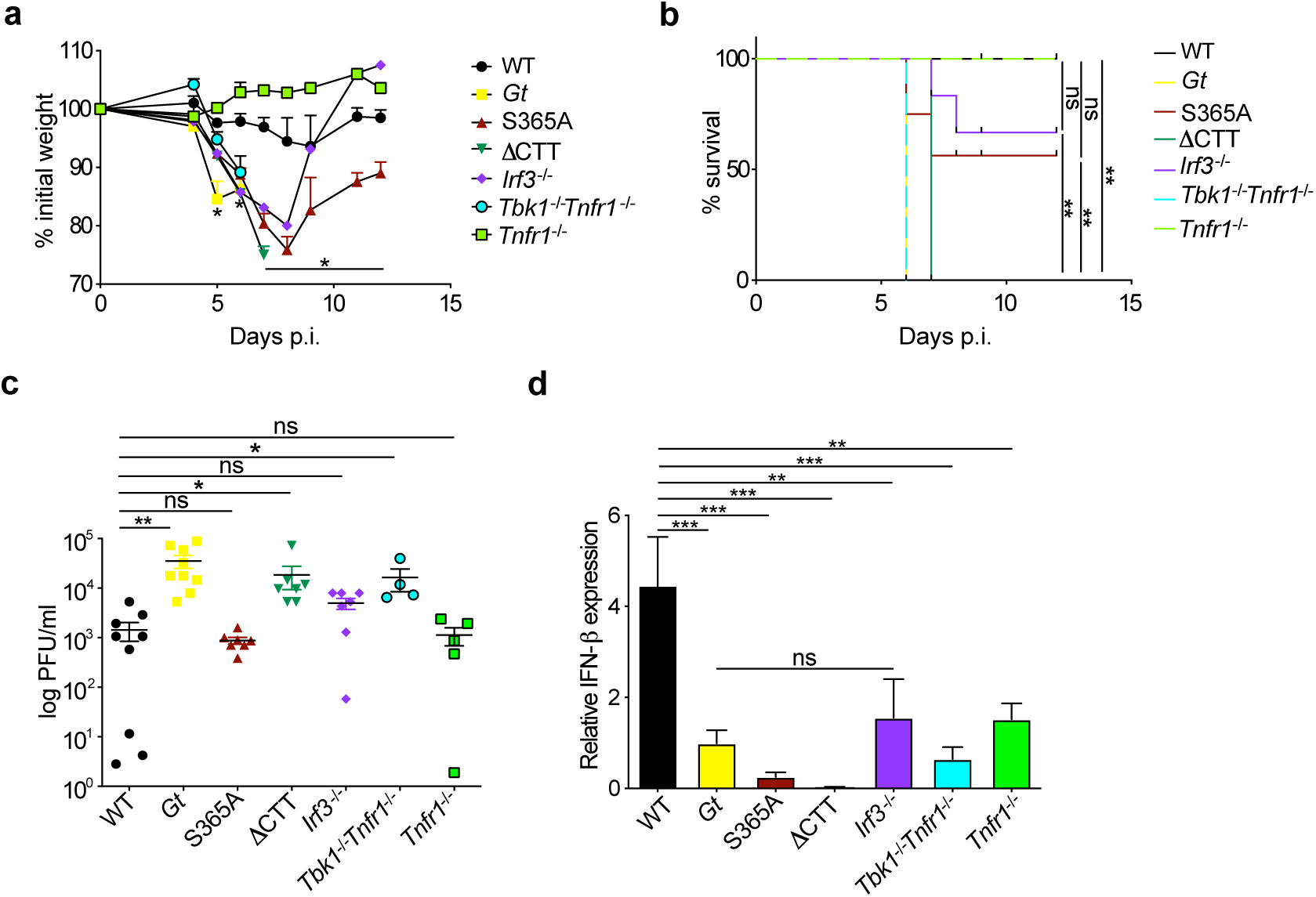
STING S365A and *Irf3*^*-/-*^ mice phenocopy resistance to HSV-1. Mice were ocular infected with 1×10^5^ PFU of HSV-1 (strain 17). **a**, Percentage of initial weight following infection. **b**, Survival of infected mice. **c**, Viral titers in the brain stem. **d**, Relative *Ifnb* expression from brain stems. Representative results of at least three independent experiments. Error bars are SEM. Analyzed with one-way ANOVA and Tukey post-test. *, p ≤ 0.05; **, p ≤ 0.005; ***, p ≤ 0.0001. ns, not significant.

## Discussion

The cGAS–STING pathway is a critical innate immune sensing pathway for the detection and elimination of DNA viruses, including HSV-1. STING activation leads to a variety of downstream antiviral responses, including IRF3-dependent induction of type I IFNs, as well as NF-kB activation and induction of autophagy responses. However, the relative contributions of these various STING-induced responses to host defense *in vivo* remains unclear. By generation and analysis of STING S365A and ΔCTT mice, we were able to investigate the role of distinct STING-dependent signaling events during HSV-1 infection. Using both a systemic and an eye infection HSV-1 model, we found that S365A mice are relatively resistant to infection, as compared to STING null *Gt* mice or to STING ΔCTT mice. STING S365A mice failed to induce type I IFNs in response to HSV-1. We therefore propose that an interferon-independent function of STING is critical during HSV-1 infection *in vivo*. IRF3 is activated downstream of STING via recruitment to phospho-S365, and IRF3 is required for type I IFN induction by STING. Interestingly, we also found that *Irf3*^*–/–*^ mice are relatively resistant to HSV-1 infection. By contrast, *Gt*, STING ΔCTT and *Tbk1*^-/-^*Tnfr1*^-/-^ mice are fully susceptible to HSV-1. TBK1 recruitment and activation by STING requires the CTT but is independent of S365. Thus, we propose that the interferon- and IRF3-independent function of STING that protects against HSV-1 is initiated upon TBK1 recruitment by the STING CTT.

Although the exact mechanism that mediates protection to HSV-1 downstream of the STING CTT and TBK1 remains to be elucidated, we propose that a strong candidate is autophagy or an autophagy-like process. Indeed, we found that STING S365A is still able to induce the autophagy-like formation of LC3 puncta (Fig. 1d-f), a process previously shown also to require TBK1 [28, 52]. Autophagy has previously been shown to be critical for control of HSV-1 [45]. However, it remains possible that an unidentified CTT–TBK1-induced response (other than, or in addition to, autophagy) is critical for STING-dependent control of HSV-1. Future studies are required to better elucidate the mechanism of STING-induced autophagy or other STING-induced responses, as there is no way at present to selectively eliminate STING-induced autophagy (or the putative autophagy-independent CTT–TBK1-dependent process). Nevertheless, our results clearly demonstrate the existence of effective S365/IRF3/interferon-independent antiviral functions for STING.

Type I IFNs are essential for control of HSV-1 [41, 43, 46-48], a result we have confirmed (Supp. Fig. S6c-d). Thus, our results suggest only that STING-induced IFN, as opposed to all sources of type I IFN, is dispensable for resistance to HSV-1. Although we observe that most type I IFN induction during HSV-1 requires STING (Fig. 4f–i), other pathways for type I IFN induction (particularly the TLR3 pathway) [54-56] have been reported and appear to provide a low but essential type I IFN response.

Autophagy has been implicated in direct antiviral defense in many neurotropic viruses infections both *in vivo* and *in vitro* [45, 57-59]. In fact, HSV-1 has evolved different mechanisms to evade autophagy [58, 60, 61], but how STING activation initiates autophagy and whether STING-induced autophagy contributes to control of HSV-1 is not clear. In addition, the involvement of TBK1 during autophagy has been a matter of discussion. Some studies show that cells lacking TBK1 can still maintain autophagy-like events (LC3 conversion, puncta formation and autophagosomes formation) [16, 62] while other evidence suggests a critical role for TBK1 in phosphorylation of selective autophagy receptors and STING autophagosomal degradation [63, 64]. Importantly, our data in primary cells suggest that TBK1 is needed for STING-mediated autophagy

One interesting feature of our results is that the STING S365A-independent protection we observe is delayed, especially in the eye infection model, and is coincident with the onset of adaptive T cell responses. Autophagy has been linked to induction of T cell responses [65-67]. Thus, one attractive possibility is that autophagy is required for antigen processing and presentation to elicit protective adaptive immune responses. Our newly generated STING mutant mice represent valuable tools to dissect this and other putative IFN-independent functions of STING *in vivo*.

## Acknowledgements

We thank members of the Vance, Barton, Stanley and Cox labs for discussions, L. Flores, P. Dietzen and R. Chavez for technical assistance, H. Nolla and A. Valeros and the Cancer Research Laboratory for flow cytometry. We thank Dr. Mary West and Dr. Pingping He of the High-Throughput Screening Facility (HTSF) at UC Berkeley. This work was performed in part in the HTSF, which provided the Perkin Elmer Opera Phenix, funded by NIH Instrument Grant S10OD021828 with the assistance of Christopher Noel. We thank Chris Bowen and Daniel Renner in the Szpara lab for assistance with viral genome sequencing and analysis. R.E.V. was supported by Investigator in the Pathogenesis of Infectious Diseases awards from the Burroughs Wellcome Fund. R.E.V. is an HHMI Investigator and is supported by NIH grants AI075039 and AI066302. The authors declare no competing interests.

## Materials and Methods

### Viruses and reagents

Dulbecco’s Modified Eagle Medium (DMEM) was obtained from Gibco and supplemented with 100 U/ml penicillin, 100 mM streptomycin and LPS-free FCS (BioWhittaker). DAPI, TRIzol, Poly I:C (all from Invitrogen) Lipofectamine 2000 (Invitrogen) were used in the experiments described below. HSV-1 (strains KOS and strain 17) was grown in Vero cells. The Vero cells used were from the lab stock. The titers of the stocks used were 8–14 × 10^9^ PFU/ml. Titers were determined by TCID50 assay on Vero cells. Both strains were used for infection of mice, while only KOS strain was used for *in vitro* stimulation.

### Mice

All mice used were specific pathogen free, maintained under a 12 h light-dark cycle (7 am to 7 pm), and given a standard chow diet (Harlan irradiated laboratory animal diet) ad libitum. Wild type C57BL/6J mice were originally obtained from the Jackson Laboratories (JAX). CRISPR/Cas9 targeting was performed by both pronucleus and cytoplasm injection of Cas9 mRNA, sgRNA, and repair template oligos into fertilized zygotes from C57BL/6J female mice (JAX, stock no. 000664), essentially as described previously[68]. STING S365A mice were generated by targeting exon 8 from STING introducing an AGT (serine) to GCC (alanine) substitution at codon 365. The sgRNA sequence was 5’ – GCTGATCCATACCACTGATG – 3’ and the repair template oligo was C*A*G*ACAAGGCTGTCCCATGCCTCAGATGAGGTCAGTGCGGAGTGGGAGA GGCTGATCCATACCGGCGATGA<u>GG</u>AGTCTTGGCTCTTGGGACAGTACGGAGG GAGGAGGTGCCACTGA*G*G*T (underlined is the PAM location). For STING ΔCTT mice, valine 340 was replaced by a premature stop codon. The sgRNA sequence was 3’ – GGAGGAAAAGAAGGACTGCT - 5’ and the repair template oligo was C*C*C*ACAGACGGAAACAGTTTCTCACTGTCTCAGGAGGTGCTCCGGCACAT TCGTCAGGAAGAAAAGGAGGA<u>GT</u>GAACCATGAATGCCCCCATGACCTCAGTG GCACCTCCTCCCTCC*G*T*A (underlined is the PAM location). The asterisks indicate phosphorothioate linkages in the first and last three nucleotides. *Irf3*^*-/-*^ mice were generated by targeting exon 6 from IRF3. The sgRNA sequence was 5’ – GAGGTGACCGCCTTCTACCG – 3’. Founder mice were genotyped as described below, and founders carrying mutations were bred one generation to C57BL/6J mice to separate modified haplotypes. Homozygous lines were generated by interbreeding heterozygotes carrying matched haplotypes. *Tbk1*^-/-^*Tnfr1*^-/-^and *Tnfr1*^-/-^ mice were described elsewhere[69].

### Preparation of gRNA transcript

DNA oligos (IDT, Coralville, NY) were heated to 95°C followed by cooling down to room temperature. The self-annealing oligo duplex was cloned into linearized T7 gRNA vector (System Biosciences, Mountain View, CA USA). The cloned sgRNA was sequence verified by DNA sequencing. Then sgRNA template for in vitro transcription (IVT) was prepared by PCR amplification of Phusion high fidelity DNA polymerase (NEB Biolabs, Ipwich, MA), the PCR mixture was cleaned up by PCR cleanup reaction (Qiagen, Hilden, Germany). The sgRNA transcripts were generated by IVT synthesis kit (System Biosciences, Palo Alto, CA). Quality of sgRNA transcripts was analyzed by NanoDrop (Thermo Fisher Scientific, Waltham, MA) and Bioanalyzer instrument (Agilent Technologies, Inc., Santa Clara, CA).

### Genotyping of STING S365A, ΔCTT and *Irf3* alleles

Exon 8 of STING and Exon 6 of *Irf3* were amplified by PCR using the following primers (all 5’ to 3’): S365A fwd: CCA ACC ATT GAA GGA AGG CTC AGT C, S365A rev: CTC ACT GTC TCA GGA GGT GCT CC; *Δ*CTT fwd: CTA GAG CCC AGA CAA GGC TGT CC, *Δ*CTT rev: CCC ACA GAC GGA AAC AGT TTC TCA C; *Irf3* fwd: AAC GTG AGT GCC AGC TGT GG, *Irf3* rev: CTT CAC AAG CTT GTC CGT CAG AAA CC. Primers were used at 200nM in a reaction with 2.5mM MgCl2 and 75μM dNTPs and 1 Unit Taq polymerase (Thermo Fisher Scientific) per reaction. Cleaned PCR products were diluted 1:16 and sequenced using Sanger sequencing (Berkeley DNA Sequencing facility).

### Cell culture

Macrophages were derived from the bone marrow of C57BL/6J or STING mutant (*Gt*, S365A or *Δ*CTT) mice. Macrophages were derived by 7 days of culture in RPMI 1640 medium supplemented with 10% serum, 100 mM streptomycin, 100 U/ml penicillin, 2 mM L-glutamine and 10% supernatant from 3T3-M-CSF cells, with feeding on day 5. Mouse primary microglia cells and astrocytes were isolated and cultured from the cerebrum of P0 pups. Neonatal cerebra were trypsinized for 20 min and filtered through a 70 μm pore size filter. Cells of 3 cerebrum were seeded on one poly-d-lysine-coated 75 cm^2^ culture flask and incubated with DMEM containing 10% FCS. The medium was replaced on day 2 after plating. Henceforth, either microglia or astrocytes were isolated. Astrocytes were isolated using the following method: after 7 days of culture, cells were shaken for 30 minutes, supernatant was aspirated, and the remaining adherent cells were predominantly astrocytes. Purity of each population was determined by FACS. Primary dissociated hippocampal cultures were prepared from postnatal day 0-1 (P0-1). Mice were euthanized using standard protocols. Briefly, bilateral hippocampi from 2-3 pups were dissected on ice and pooled together. The tissue was dissociated using 34.4ug/ml papain in dissociation media (HBSS Ca2+, Mg2+ free, 1mM sodium pyruvate, 0.1% D-glucose, 10mM HEPES buffer) and incubated for 3 min at 37° C. The papain was neutralized by incubation in trypsin inhibitor (1mg/ml in dissociation media) at 37°C for 4 min. After incubation, the dissociation media was carefully removed and the tissue was gently triturated, manually, in plating media (MEM, 10% FBS, 0.45% D-Glucose, 1mM sodium pyruvate, 1mM L-glutamine). Cell density was counted using a TC10 Automated cell counter (Biorad). For western blot experiments, 2.2-2.5 × 10^5^ cells were plated onto 24-well plates pre-coated with Poly-D-Lysine (PDL) (Corning) in 500ul of plating media. After 3 hours, plating media was removed and 800ul maintenance media (Neurobasal media (GIBCO) with 2mM glutamine, pen/strep, and B-27 supplement (GIBCO)) was added per well. After 4 days in vitro 1uM cytosine arabinoside (Sigma) was added to prevent glial proliferation. Neurons were maintained in maintenance media for 14 days with partial media changes every 4 days. For immunofluorescence, 2 × 10^3^ cells were plated in pre-coated 96 well plates (CellCarrier-96 Ultra Microplates, black, PerkinElmer) following the same procedure.

### Murine *M. tuberculosis* infections

*M. tuberculosis* strain Erdman (gift of S.A. Stanley) was used for all infections. Frozen stocks of this wild-type strain were made from a single culture and used for all experiments. Cultures for infection were grown in Middlebrook 7H9 liquid medium supplemented with 10% albumin-dextrose-saline, 0.4% glycerol and 0.05% Tween-80 for five days at 37°C. Mice were aerosol infected using an inhalation exposure system (Glas-Col, Terre Haute, IN). A total of 9 ml of culture was loaded into the nebulizer calibrated to deliver ∼400 bacteria per mouse as measured by colony forming units (CFUs) in the lungs 1 day following infection (data not shown). Mice were sacrificed at various days post-infection as indicated in the figure legends to measure CFUs and/or cytokines. All lung lobes were homogenized in PBS plus 0.05% Tween-80 or processed for cytokines (see below), and serial dilutions were plated on 7H11 plates supplemented with 10% oleic acid, albumin, dextrose, catalase (OADC) and 0.5% glycerol. CFUs were counted 21-25 days after plating.

### Cytokine measurements

Cell-free lung homogenates from *M. tuberculosis* infected mice were generated as previously described[70]. Briefly, lungs were dissociated through 100 μm Falcon cell strainers in sterile PBS with 1% FBS and Pierce Protease Inhibitor EDTA-free (Thermo Fisher). An aliquot was removed for measuring CFU by plating as described above. Cells and debris were then removed by first a low-speed centrifugation (approximately 300×g) then a high-speed centrifugation (approximately 2000×g) and the resulting cell-free homogenate was filtered twice with 0.2 μm filters to remove all *M. tuberculosis* for work outside of BSL3. All homogenates were aliquoted, flash-frozen in liquid nitrogen and stored at –80°C. Each aliquot was thawed a maximum of twice to avoid potential artifacts due to repeated freeze-thaw cycles. All cytokines were measured using Cytometric Bead Assay (BD Biosciences) according to manufacturer protocols. TNF-α from DMXAA and LPS stimulated mice was also measured by CBA. Results were collected using BD LSRFortessa (BD Biosciences) and analyzed using GraphPad Prism v6.0c. TNF-α from primary macrophages supernatant was measured by ELISA.

## Murine HSV-1 infection models

### Intravenous infection

Age and sex matched (7–10-week old) mice were warmed under a lamp for venous dilation and inoculated with 1 × 10^6^ PFU HSV-1 (KOS strain) in 200μl of PBS or mock infected with PBS only.

### Ocular infection

Age and sex matched (7–10-week old) mice, were anaesthetized with intraperitoneal (i.p.) injection of ketamine (100 mg/kg body weight) and xylazine (10 mg/kg body weight). Corneas were scarified using a 25G needle and mice were either inoculated with 1 × 10^5^ PFU HSV-1 (strain 17) in 5 μl, or mock infected with 5 μl of PBS. Eyewash was collected by gently proptosing each eye and wiping a sterile cotton swab around the eye in a circular motion. The swabs were placed in 0.5 ml of DMEM medium and stored at −80 °C until the titer was determined. Whole brains, brain stems, spinal cords and livers were frozen immediately at −80 °C. Tissues were homogenized with tissue homogenizer (Polytron PT 2500 E) for 2 min at frequency 10. Tissues were used for RNA isolation with TRIzol or used for virus titration.

### Scoring and tissue harvest

Mice were scored for disease, weighed at the indicated times post infection and euthanized at the specified times post infection for tissue harvesting or once they met end point criteria. The scoring was performed as blinded study, largely following previous descriptions by others[51] with the following minor modifications: symptoms related to neurological disease named body condition score (BCS) (0: normal, healthy 1: hunched, 2: uncoordinated, lethargic, mild paralysis, 3: unresponsive/no movement, complete paralysis).

### Infection and cell stimulations (transfections)

For infections, bone marrow derived macrophages from C57BL/6J mice were plated at 1-2×10^6^ cells/well. The next day they were stimulated with cyclic dinucleotides c-di-GMP, 2’3’cGAMP, Sendai virus (SeV) and poly I:C. Cells were transfected using Lipofectamine 2000 (LF2000; Invitrogen) according to the manufacturer’s protocol. All cyclic dinucleotides nucleic acid stimulants were mixed with LF2000 at a ratio of 1 μl LF2000/1 μg nucleic acid, incubated at room temperature for 20–30 min, and added to cells at a final concentration of 4 μg/ml (6-well plates). For Sendai Virus, cells were infected at 150 hemagglutination units (HAU)/ml. For poly I:C, 2 mg/ml of the stock solution was heated at 50°C for 10 min and cooled to room temperature before mixing with LF2000. Transfection experiments were done for 6 h, unless otherwise stated in the figures.

### Immunoblotting

BMMs were seeded at a density of 1×10^6^ cells per well in 6 well tissue culture plates and transfected the next day using Lipofectamine 2000 (Invitrogen) according to the manufacturer’s instruction. Cells were lysed at indicated time post transfection with radioimmunoprecipitation assay (RIPA) buffer supplemented with 2 mM NaVO3, 50 mM b-Glycerophosphate, 50 mM NaF, 2 mM PMSF, and Complete Mini EDTA-free Protease Inhibitor (Roche). Proteins separated with denaturing PAGE and transferred to Immobilon-FL PVDF membranes (Millipore). Membranes were blocked with Li-Cor Odyssey blocking buffer. Primary antibodies were added and incubated overnight. Primary antibodies used were: anti-TBK1 (D1B4) (#3504), anti-phospho-TBK1/NAK (Ser172) (D52C2) (#5483), anti-STING (D2P2F) (#13647), anti-phospho-STING (Ser366) (D7C3S) (#19781), anti-phospho-IRF3 (Ser396) (4D4G) (#4947), all purchased from Cell Signaling Technologies. Anti-IRF3 (EP2419Y) (#ab76409) was from Abcam. Secondary anti-rabbit IgG was conjugated to Alexa Fluor-680 (Invitrogen). Immunoblots were imaged using a Li-Cor fluorimeter.

### Quantitative PCR

Stimulated cells were overlayed with TRIzol (Invitrogen) and stored. RNA was isolated according to the manufacturer’s protocol and was treated with RQ1 RNase-free DNase (Promega). 0.5 μg RNA was reverse transcribed with Superscript III (Invitrogen). SYBRGreen dye (ThermoFisher Scientific) was used for quantitative PCR assays and analyzed with a real-time PCR system (StepOnePlus; Applied Biosystems).

All gene expression values were normalized to *Rps17* (mouse) levels for each sample. The following primer sequences were used: mouse *Ifnb*, (forward) 5′-ATAAGCAGCTCCAGCTCCAA-3′ and (reverse) 5′-CTGTCTGCTGGTGGAGTTCA-3′; mouse *Rps17*, (forward) 5′-CGCCATTATCCCCAGCAAG-3′ and (reverse) 5′-TGTCGGGATCCACCTCAATG-3′; mouse *Viperin*, (forward) 5′-TTGGGCAAGCTTGTGAGATTC-3′ and (reverse) 5′-TGAACCATCTCTCCTGGATAAGG-3′; mouse *TNF*, (forward) 5′-TCTTCTCATTCCTGCTTG TGG-3′ and (reverse) 5′-GGTCTGGGCCATAGAACTGA-3′; mouse *IL-6*, (forward) 5′-GCTACCAAACTGGATATAATCAGGA-3′ and (reverse) 5′-CCAGGTAGCTATGGTACTCCAGAA-3′.

### Immunofluorescence and high-content imaging

Bone marrow derived macrophages were transfected with 0.2 ug of Cy3-labeled DNA for 6 hours. Cells were washed with PBS, fixed in 4% paraformaldehyde and ice-cold methanol. Cells were washed 3x with PBS and blocked and permeabilized with 2% BSA and 0.3% Triton X100. LC3 puncta staining was performed using mouse monoclonal antibody (Nanotools, catalog #0260-100/LC3-2G6 at 1:400, RT) for 3hours, followed by secondary goat anti-mouse IgG labeled with Alexa Fluor 488 (Life Technologies at 1:4000, RT) for 1 hour. Nuclei were stained with DAPI. For imaging, cells in 96-well plates were imaged using an Opera Phenix (Perkin Elmer) at RT, using a × 40 1.1 NA water immersion lens (Zeiss). Images were exported to Harmony High-Content Imaging and Analysis Software and automated colocalization measurements were performed with the Perkin Elmer Harmony software package. A pipeline was created to measure colocalization of Cy3-labeled DNA and LC3. Quantification was performed using data collected from 16 fields per well in 96-well format. Data was then analyzed in Prism using one-way ANOVA analysis.

### Flow cytometry

Single suspensions were prepared from each experimental group using a modified protocol as described[71]. To analyze tetramer positive cells, cell suspensions were stained with the following cell surface antibodies: CD3e (clone 145-2C11, BD Horizon), CD8a (clone 53-6.7, Biolegend), CD45 (clone 30-F11, eBioscience), CD44 (clone IM7, eBioscience), CD11b (clone M1/70, eBioscience), MHCII I-A/I-E (cloneM5/114.15.2, Biolegend), CD19 (clone eBio1D3, eBioscience), CD45R (B220) (clone RA3-6B2, Invitrogen), Ly6G (Gr-1) (clone 1A8-Ly6g, eBioscience). Samples were acquired on a FACS X20 Fortessa (BD Bioscience) and analyzed with FlowJo software (TreeStar).

### Statistical analysis

All data were analyzed with one-way ANOVA test and Tukey post-test unless otherwise noted and survival data were analyzed with Log-rank (Mantel-Cox) test. Both tests were run using GraphPad Prism 6. *, p ≤ 0.05; **, p ≤ 0.005, ***, p ≤ 0.0001. All errors bars are SEM and all center bars indicate means.

## Supplementary figures

**Fig. S1.**
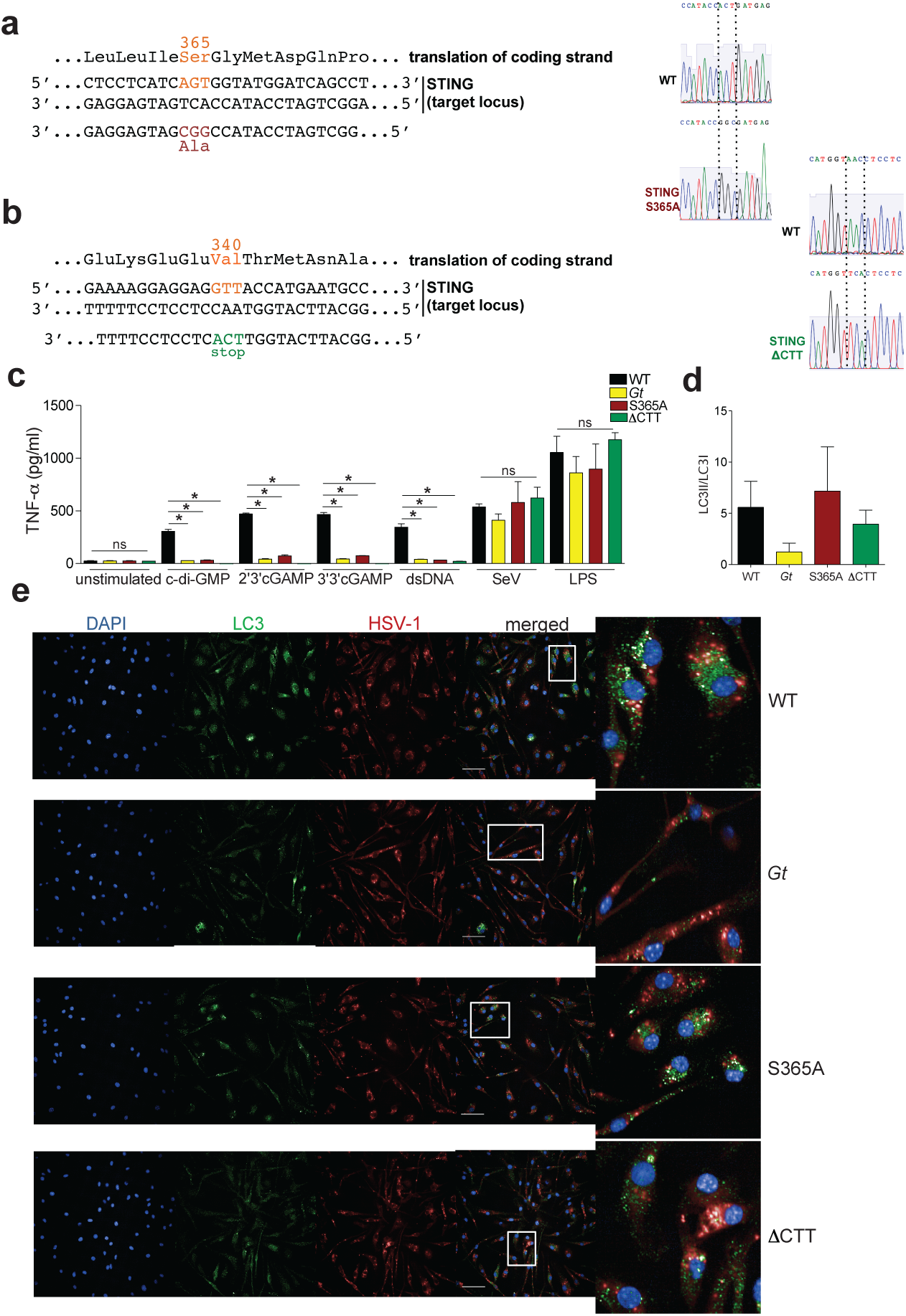
(related to figure 1). **a**, Creation of STING S365A and **b**, ΔCTT mice using CRISPR/Cas9. **c**, Bone marrow derived macrophages were stimulated for 6h and TNF-α was measured on the supernatant. **d**, Quantification of Fig.1d using LC3II/LC3I ratio. **e**, Colocalization of DNA and LC3 is increased in WT and S365A cells. Fluorescence images of primary macrophages transfected for 6h with Cy3-labeled DNA and LC3. Images were analyzed by an automated pipeline created on Perkin Elmer Harmony software for colocalization quantification (for more details refer to Methods). Scale bars are 50 μm.

**Fig. S2.**
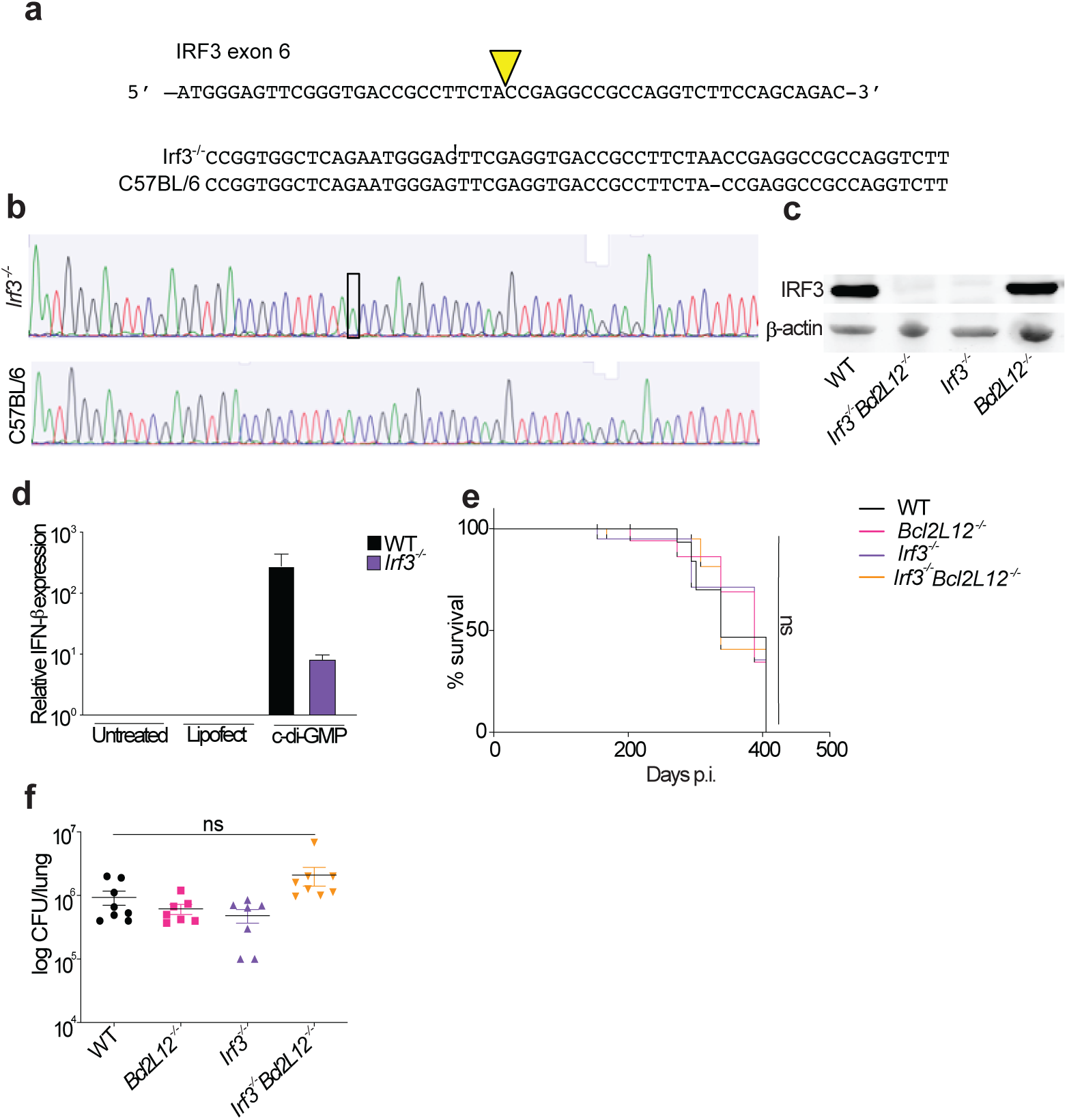
(related to figure 2). Creation of IRF3 deficient mice using CRISPR/Cas9. **a**, CRISPR/Cas9 targeting strategy for IRF3. **b**, Sequencing of the targeted locus resulting in *Irf3*^*-/-*^ mutation. **c**, Immunoblot of MEFs for IRF3. **d**, Primary macrophages were transfected with c-di-GMP for 6h and relative expression of *Ifnb* was analyzed. **e**, Mice were aerosol infected with 400 CFU dose of *M. tuberculosis* (Erdman strain). Survival of infected mice. **f**, Bacterial burden from lungs at 21 days post infection. All mice except C57BL/6J WT were bred in-house. Representative results of four independent experiments. Error bars are SEM. Analyzed with one-way ANOVA and Tukey post-test. ns, not significant.

**Fig. S3.**
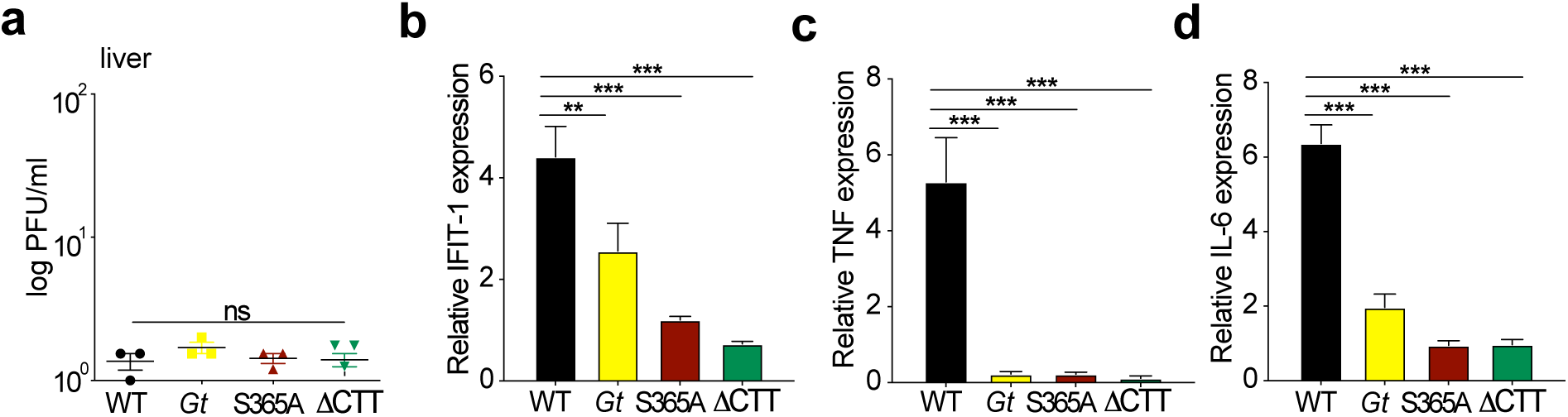
(related to figure 3). Mice were intravenously infected with 1×10^6^ PFU of HSV-1 (KOS strain). **a**, Viral titers in the liver at 6 days p.i. **b**, Relative expression of IFIT-1 **c**, TNF and **d**, IL-6 from brains at 3 days p.i. All mice except C57BL/6J WT were bred in-house. Representative results of five independent experiments. Error bars are SEM. Analyzed with one-way ANOVA and Tukey post-test. **, p ≤ 0.005; ***, p ≤ 0.0001. ns, not significant.

**Fig. S4.**
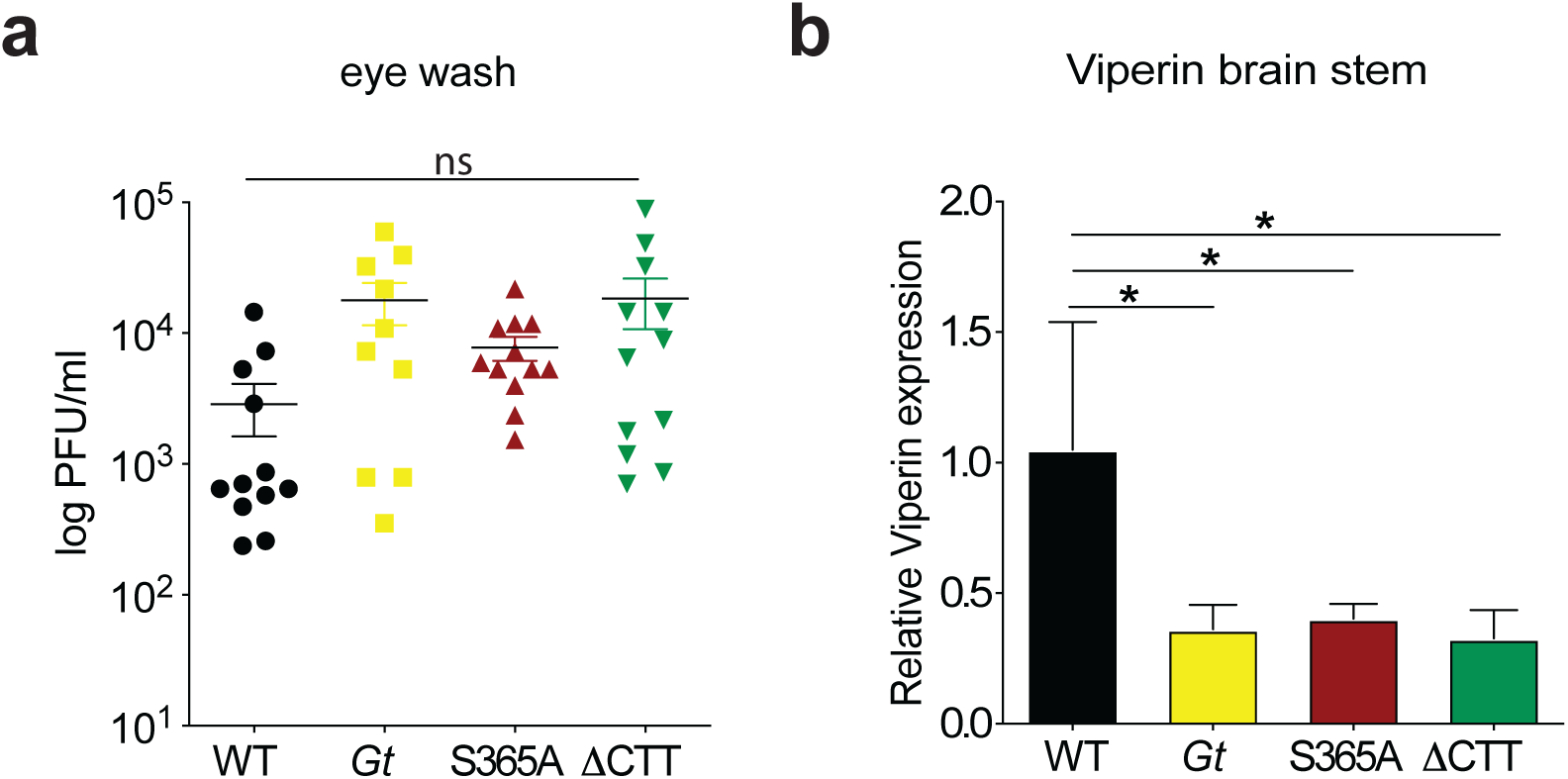
(related to figure 4). Mice were ocular infected with 1×10^5^ PFU of HSV-1 (strain 17). **a**,Viral titers from eyes washed at 2 days p.i.. **b**, Relative expression of *viperin*. All mice except C57BL/6J WT were bred in-house. Representative results of three independent experiments. Error bars are SEM. Analyzed with one-way ANOVA and Tukey post-test. ns, not significant.

**Fig. S5.**
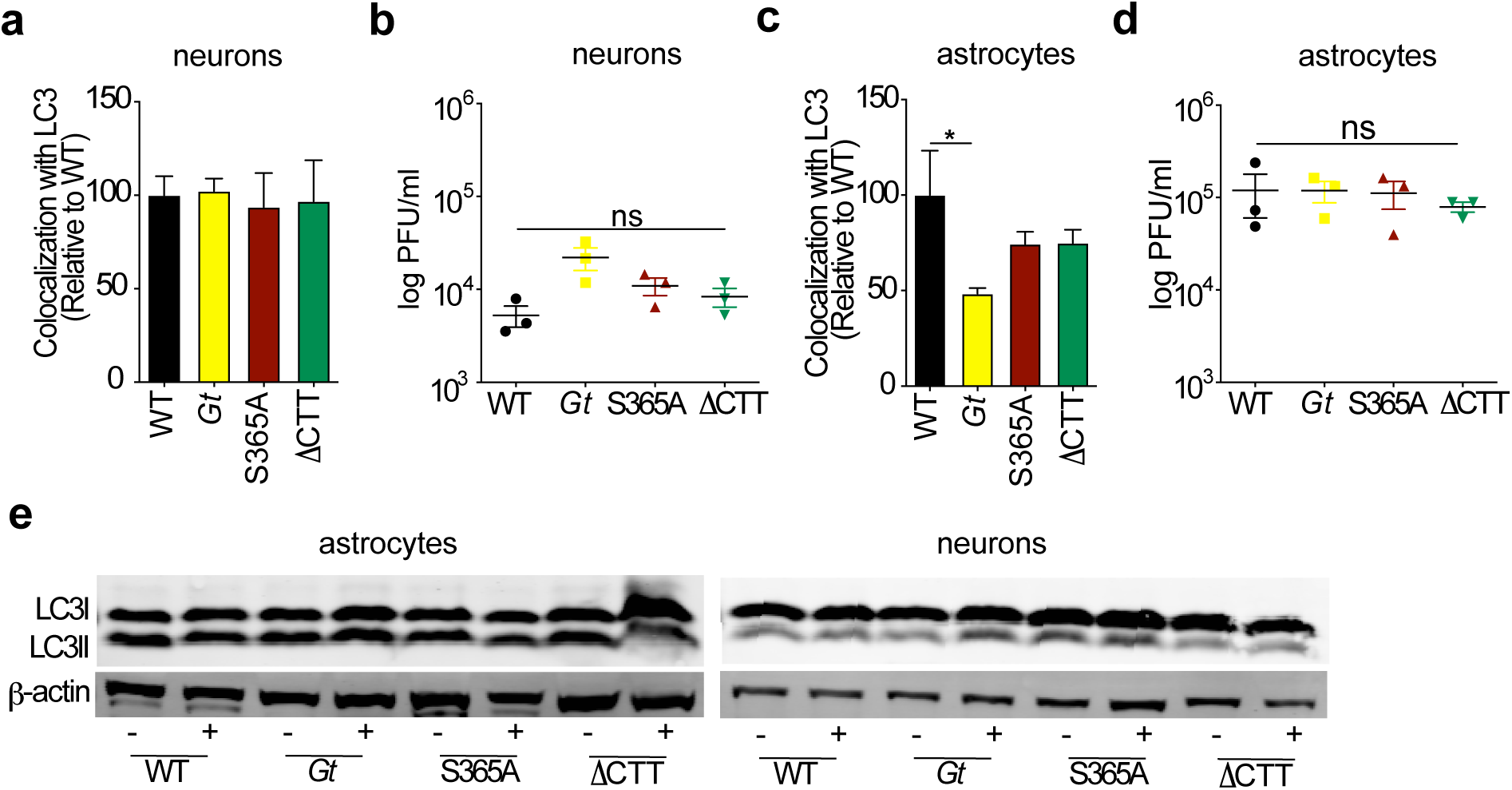
(related to figure 4). Brain cells (neurons and astrocytes) were harvested from P0 pups and infected with HSV-1 (KOS strain) at a MOI 1 for 6h and later were stained for LC3 and HSV-1. **a**, Quantification of colocalization of LC3-HSV-1 in neurons was performed and **b**, Viral titers from supernatants were collected 48h later and quantified by TCID50 assay. **c-d**, Same as a-b, in astrocytes. **e**, Cell lysates were collected at 4h post-infection and immunoblot for LC3 and β-actin was performed. Representative results from two independent experiments. Error bars are SEM. Analyzed with one-way ANOVA and Tukey post-test. *, p ≤ 0.05. ns, not significant.

**Fig. S6.**
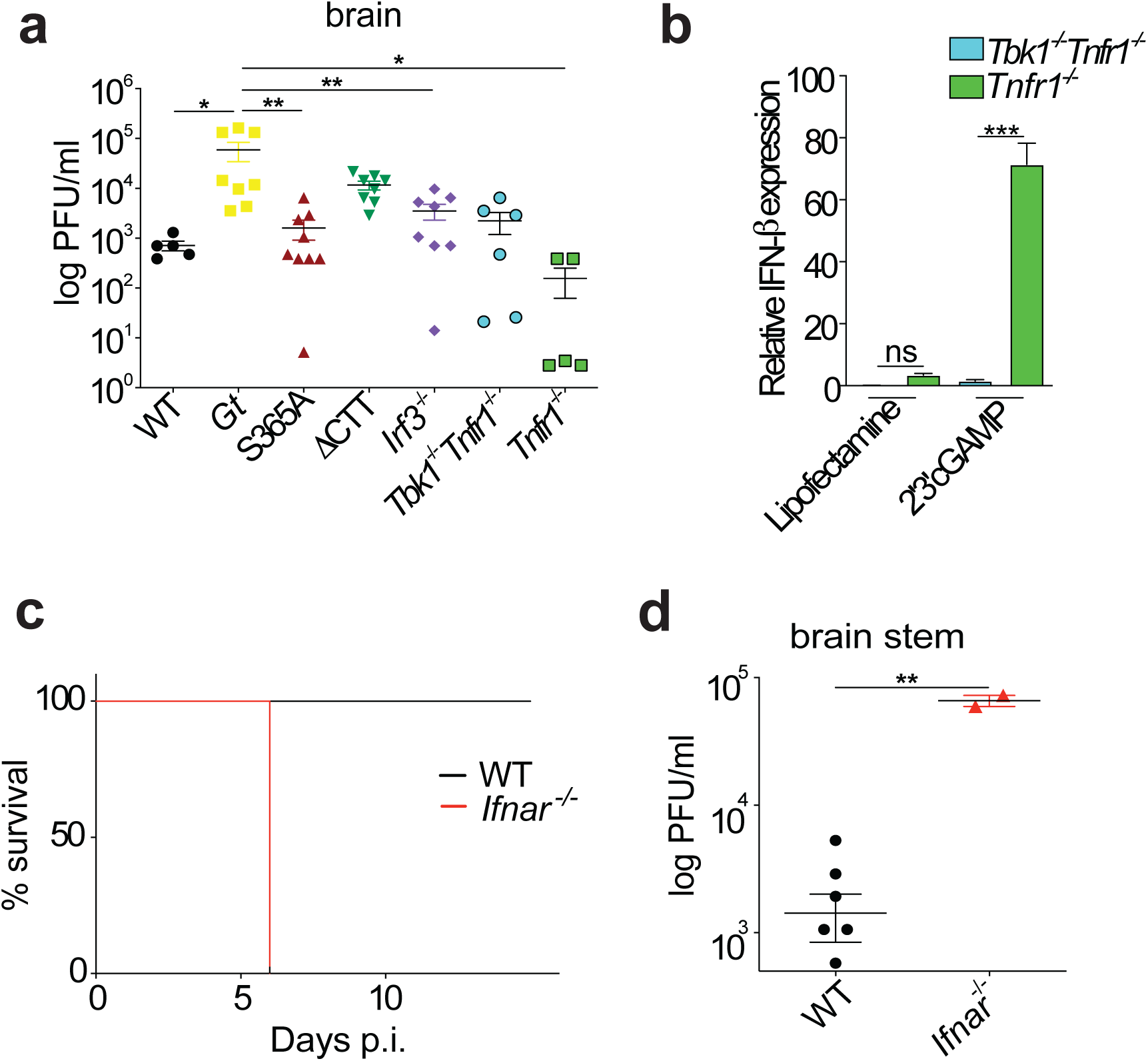
(related to figure 5). **a**, Mice were ocular infected with 1×10^5^ PFU of HSV-1 (strain 17) and viral titers measured in the brain 6 days p.i. **b**, BMDMs were transfected with 2′3′cGAMP for 6h and relative expression of *Ifnb* was analyzed. **c**, Mice were ocular infected with 1×10^5^ PFU of HSV-1 (strain 17) and survival rate and **d**, viral titers measured in the brain stem 6 days p.i. All mice except C57BL/6J WT were bred in-house. Representative results of two independent experiments. Error bars are SEM. Analyzed with one-way ANOVA and Tukey post-test. *, p ≤ 0.05; **, p ≤ 0.005. ns, not significant.

